# Juvenile-mimicry explains adult-juvenile resemblance in swallows and martins (Aves: Hirundinidae)

**DOI:** 10.64898/2026.05.26.728048

**Authors:** Masaru Hasegawa

**Author notes:** Corresponding author: Dr Masaru Hasegawa, Nara University of Education, 630-8528 Takabatake-Cho Nara City, Nara Prefecture, Japan.

## Abstract

A similar phenotype exhibited by both adults and juveniles is often considered a self-evident default state due to shared genes and similar ecological niches, and thus the function of adult-juvenile resemblance is rarely addressed. An adaptive explanation for adult-juvenile resemblance is that adults mimic juveniles to attract mates by exploiting their parental care behavior and to avoid agonistic intrasexual combat from rivals that tolerate juveniles (i.e., the juvenile-mimicry hypothesis). Using a phylogenetic comparative approach, we tested the juvenile-mimicry hypothesis in aerial foragers, swallows and martins (Aves: Hirundinidae), in which adults and juveniles frequently encounter one another in their open habitat. We predicted that, if adults mimic juveniles, adult-juvenile resemblance should be enhanced in species with many young (i.e., a large number of models in relation to mimics) as well as species with a few young (i.e., a default state with limited intensity of sexual selection). This prediction was confirmed by a quadratic relationship between number of juveniles and adult-juvenile resemblance. In addition, as predicted under the juvenile-mimicry hypothesis, adult-juvenile resemblance was enhanced in species with multiple broods, in which juvenile-mimicry would be particularly effective due to the mating period followed by juvenile production. The observed pattern could not be explained by sexual selection for male ornamentation alone (i.e., with no juvenile-mimicry) even when considering the cost of ornamentation. An alternative explanation that juveniles mimic adults is also unlikely, as the situation favors the opposite pattern. The current study therefore supports the juvenile-mimicry hypothesis, indicating an adaptive function of adult-juvenile resemblance.

## Introduction

Adult animals often possess several conspicuous secondary sexual characteristics and thus can easily be distinguished from juveniles (Andersson 1994). Such characteristics are considered to have evolved by sexual selection, and in fact a number of empirical studies support this explanation by demonstrating evidence of sexual selection on these characteristics (e.g., Hill and McGraw 2006; Tazzyman et al. 2014 for recent reviews). In other species, however, adults closely resemble juveniles with no apparent secondary sexual characteristics, which is often considered a self-evident “default” state due to shared genes and seemingly similar ecological niches, with little sexual selection on the adult phenotype (the “null hypothesis” hereafter). For this reason, secondary sexual characteristics and their sexual dimorphisms, opposite to the default, are frequently used as an index of sexual selection (Kraaijeveld et al. 2011). Adult-juvenile resemblance itself is therefore thought to be functionless and is rarely addressed. However, this explanation ignores the fact that adults but not juveniles participate in reproduction, including mating and parental care, and thus ecological niches of adults and juveniles are in fact differentiated. Adult phenotypes are better adapted to reproduction, radically counteracting adult-juvenile resemblance.

An alternative, adaptive explanation of adult-juvenile resemblance is given by the “juvenile-mimicry hypothesis”: adults mimic nonreproductive individuals (i.e., juveniles) as “models” to avoid aggression from “signal receivers,” conspecific rivals (e.g., Burley 1981; Foster 1987; reviewed in Senar 2006; Rainey and Grether 2007; Hawkins et al. 2012). In fact, the juvenile phenotype can have adaptive functions to avoid agonistic interactions with conspecific adults that are competitively superior to juveniles (e.g., Ligon et al. 2009; reviewed in Senar 2006), and thus a juvenile-like phenotype would be beneficial for adults as well to avoid aggression (reviewed in Hawkins et al. 2012). The juvenile phenotype can reduce adult aggression even in species with little parental care (e.g., Fresnillo et al. 2015), perhaps to avoid wasting time and energy on non-rival conspecifics (i.e., juveniles), but juvenile mimicry would be particularly beneficial in species with parental care in which adults provide parental care to their young and thus suppress the agonistic reaction to them (Christy 1995; Arnqvist 1998; see below).

In species with parental care, adults are attracted to traits that resemble those of offspring, i.e., young-like traits, and thus such traits can be used as a sensory trap to attract potential mates by exploiting their parental behavior (i.e., attraction to young-typical traits; e.g., Stålhandske 2002; Amcoff et al. 2013; Hasegawa et al. 2013). Sensory traps (i.e., young-like traits, here) can be maintained in the population even if they are costly to signal receivers when strong selection favors their response to the model (i.e., young traits; e.g., see Christy 1995; Arnqvist 1998; note that initially maladaptive response can be beneficial; Marcias-Garcia and Ramirez 2005). Although these reports have often focused on female responses, not only females but also males are “trapped” by mimetic signals and suppress their aggressive reactions (Hasegawa 2021). Therefore, adult-juvenile resemblance itself has dual function in sexual selection at least in some circumstances and hence interspecific variation in “secondary sexual characteristics (or its inverse, adult-juvenile resemblance)” might be explained by sexual selection for juvenile-mimicry. A macroevolutionary study on adult-juvenile resemblance supported this hypothesis, because cuckoo species with parental care behavior resemble their young more than brood-parasitic cuckoos (Hasegawa and Arai 2018a), but such an approach still suffers from confounding factors between the two groups (e.g., the presence or absence of brood-parasitism). A better approach would be to focus on species with parental care with varying degrees of adult-juvenile resemblance.

Swallows and martins are suitable study systems for this purpose, because they show a variety of plumage phenotypes ranging from a high degree of adult-juvenile resemblance (e.g., the white-thighed swallow *Neochelidon tibialis*, in which adults and juveniles are very similar) to some differences between adult and juvenile phenotypes (e.g., the barn swallow *Hirundo rustica*, in which juveniles have duller and browner plumage with shorter tails and paler rufous areas than adults; Turner and Rose 1994). Like other birds, adult and juvenile hirundines have similar body sizes, and thus juvenile-mimicry can be easier than in taxa with marked differences in size between adults and juveniles (e.g., fish, reptiles; Fresnillo et al. 2015). Moreover, hirundines typically share a very similar ecology, including aerial foraging, annual molt, social monogamy, and biparental provisioning (Turner and Rose 1994), which is suited to study adult-juvenile resemblance with limited confounding effects of ecological diversification. Aerial foraging, in particular, provides a suitable opportunity to study adult-juvenile resemblance, as there are frequent encounters between adults and juveniles in the open habitat with no place to hide (Turner and Rose 1994), meaning that their appearance cannot be free from selection via intraspecific interactions. Furthermore, recent experiments with barn swallows have demonstrated that nestling-like vocalizations can function for attracting social mates and avoiding aggression from neighboring rivals (Hasegawa et al. 2013; Hasegawa and Arai 2016; Hasegawa 2021), indicating that juvenile-mimicry would be functional in inter- and intrasexual selection in at least some hirundines (also see Turner and Rose 1994). In fact, Turner (2006) pointed out aggression avoidance of juveniles (and juvenile-like plumage coloration of yearlings, though these are thought to be “female-mimicry”).

Here we tested the juvenile-mimicry hypothesis in the context of sexual selection using a phylogenetic comparative analysis of swallows and martins (Aves: Hirundinidae). The juvenile-mimicry hypothesis predicts that adult-juvenile plumage resemblance should be enhanced in species with large broods, which can have a large number of models (i.e., many juveniles in hirundines, in which fledging success can be high: Turner and Rose 1994). In other words, as the number of juveniles increases, the direction of sexual selection would bias more toward juvenile-like phenotype rather than the opposite (i.e., toward pronounced secondary sexual characteristics) due to selection favoring more-juvenile like adults. This changing direction of sexual selection, or increased “weight” of juvenile-like phenotype, should be distinguished from changes in the overall intensity of sexual selection, which can also increase with the number of juveniles due to the increased variance in offspring numbers (e.g., variance is proportional to mean values when it follows Poisson distribution, and this is intuitively straightforward as difference in seasonal reproductive success of paired and unpaired individuals increases with mean number of offspring). Then, assuming the presence of sexual selection for secondary sexual characteristics (see the first paragraph), we can predict a quadratic relationship between the number of juveniles, i.e., index of opportunity for sexual selection via social offspring, and the measure of secondary sexual characteristics (i.e., adult-juvenile plumage discrepancy, here) under the juvenile-mimicry hypothesis (Fig. 1, right panels). That is, adults would closely resemble juveniles in species with many young (i.e., a large number of “models” in relation to “mimics”) as well as species with a few young (i.e., a default state with limited intensity of sexual selection). Such a quadratic relationship is not predicted by sexual selection for secondary sexual characteristics alone, even when assuming the increasing cost of secondary sexual characteristics, which can simply offset the increasing benefits, with the number of juveniles (see Fig. 1 for an explanation; also see Sterns 1992; Fitzpatrick et al. 1995; Badyaev 1997 for cost of ornamentation in the context of rearing young).

**Fig. 1.**
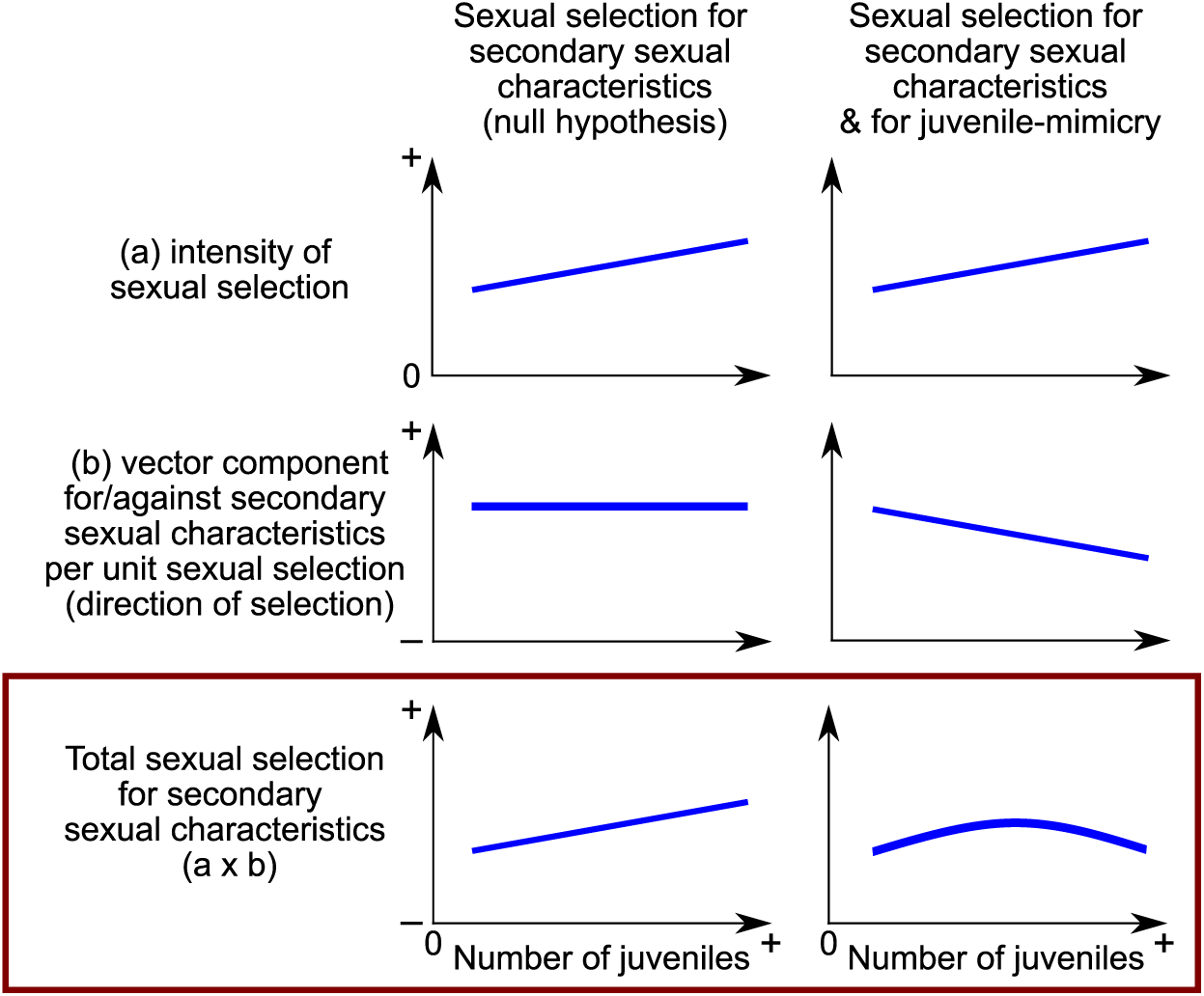
Expectations of alternative hypotheses. Sexual selection for secondary sexual characteristics without juvenile-mimicry (null hypothesis, here: left panels), in which the juvenile-like phenotype is regarded as a default state, would favor the increased expression of overall secondary sexual characteristics when opportunities for sexual selection is high. In this case, overall expression of secondary sexual characteristics increases with increasing number of juveniles, because intensity of sexual selection would increase with the mean number (and hence its variance as well) of juveniles (see text for details). This prediction can be slightly changed when secondary sexual characteristics impose a substantial rearing cost, in which the expression of secondary sexual characteristics can decrease (rather than increase) with increasing number of juveniles due to the increased cost of rearing juveniles, depending on the relative intensity of sexual and viability selection. In contrast to the null hypothesis (and its variant with respect to the increased cost of secondary sexual characteristics with the number of juveniles, which can change the gradient), sexual selection for juvenile-mimicry is unique as it predicts a quadratic relationship with the number of juveniles would be uniquely found for adult-juvenile plumage discrepancy (right panels), as juvenile-like phenotype, i.e., the inverse of secondary sexual characteristics, is relatively more beneficial when models are more abundant (i.e., when number of juveniles increases; right panels). We focus here solely on sexual selection on secondary sexual characteristics, as all other sexual selections orthogonal to the sexual selection on secondary sexual characteristics are irrelevant. Feedback of viability cost of secondary sexual characteristics on the sexual selection on secondary sexual characteristics is not straightforward and thus not taken into account, because the function of sexual selection for secondary sexual characteristics depends on differential cost (rather than mean cost) of secondary sexual characteristics (i.e., signal receivers may still favor “costly” traits if they impose higher cost on low-quality individuals, even if cost of the traits on average is heightened; note that the benefits of signal senders using the sexual traits depend on the intensity of sexual selection, further complicating potential feedback effects)

These predictions, however, are based on simple expectations where viability selection and sexual selection, in which all components are assumed to be a linear function, additively determine total selection for (and hence expression of) secondary sexual characteristics. Changing assumptions (e.g., exponential cost rather than linear cost; multiplicative, rather than additive, determination of total selection; e.g., Fromhage et al. 2022) can affect predictions: for example, multiplicative determination of total selection predicts a quadratic relationship between number of young (n) and the expression of secondary sexual characteristics, as viability selection (denoted as a_v_n + b_v_, in which a_v_ and b_v_ is the slope and intercept, respectively) multiplied by sexual selection (likewise, a_s_n + b_s_) provides a quadratic function [a_v_a_s_n^2^ + (a_v_b_s_ + a_s_b_v_)n + b_v_b_s_]. For this reason, we also analyzed male-female plumage discrepancy (i.e., sexual plumage dimorphism), a well-known measure of sexual selection for costly traits. Sexual selection theory predicts that sexual plumage dimorphism would be expected in costly sexual traits under stronger sexual selection for such male traits and/or stronger viability selection against homologous female traits (e.g., see Dale et al. 2015). Thus, if costliness of secondary sexual characteristics drives the observed pattern of adult-juvenile plumage discrepancy (e.g., via an exponential cost function or multiplicative effect of sexual and viability selection; see above), we can predict that sexual plumage dimorphism increases with the number of juveniles. Or, if overall intensity of sexual selection is enhanced at the intermediate number of juveniles for unknown reason, we can expect the quadratic relationship between number of juveniles and secondary sexual characteristics (as well as adult-juvenile plumage discrepancy). We evaluated absolute measure of male plumage appearance, which is obtained from Dale et al. (2015). Given that juvenile-mimicry selects on adult plumage appearance “relative” to juveniles, a quadratic relationship with the number of juveniles would be uniquely found for adult-juvenile plumage discrepancy (i.e., not for absolute male plumage appearance) under the juvenile-mimicry hypothesis.

In addition, a recent phylogenetic comparative study showed that incubation type can be used as a proxy for extrapair mating opportunities, which can increase sexual selection on plumage ornamentation and sperm quality (Hasegawa and Arai 2020b, 2022): species with female-only incubation have more than three times as many extrapair mating opportunities as species with biparental incubation, perhaps because incubation effort and extrapair mating effort cannot be performed simultaneously. Therefore, we also investigated the possible interacting effect of incubation type and number of juveniles on secondary sexual characteristics (i.e., adult-juvenile plumage discrepancy). Because sexual selection for seemingly juvenile-mimicry is used in social mating (but not in extrapair mating) at least in the barn swallow (Turner 2006; Hasegawa et al. 2013), we expected that changing direction of sexual selection should be more evident in species with low opportunities of extrapair mating (i.e., in which social mating should be relatively important). Lastly, in addition to the abovementioned predictions on spatial abundance of models and mimics, we can also predict that juvenile-mimicry should be most effective when the mating period, in which sexual selection mainly occur, overlaps with the juvenile period (i.e., when models and mimics certainly co-exist at the same time). In fact, Hasegawa et al. (2013) discussed that mimicry of young would have originated in situation where males court females soon after the nestling period. In this sense, we can predict that species with multiple broods, i.e., those having the mating period after juvenile production, would have higher adult-juvenile similarity, though this is not the sole opportunity for adults encounter with juvenile during the mating season (e.g., asynchronous breeding and re-clutches after failed reproduction increase inter-individual variation in the timing of the mating period, though these data are difficult to obtain).

We discuss the observed pattern from the evolutionary perspective. Although we focused on the number of juveniles per breeding pair (i.e., relative abundance of “model” in relation to potential “mimic”), one may argue that breeding density is another factor potentially enhancing the benefits of juvenile mimicry, as the number of juveniles increase with population density. However, the number of adults increases with increasing density (i.e., the adult:juvenile ratio does not simply increase with population density); thus, we cannot easily make predictions for the adult-juvenile discrepancy in relation to breeding density. This contrasts with our prediction based on the number of young per breeding pairs (see above), because mimic (i.e., juvenile-like adults) can spread in the population when their relative abundance to model (i.e., juveniles) is low, as is the case in interspecific mimicry (e.g., Lindström et al. 1997).

## Methods

### Data collection

As in previous studies (e.g., Hasegawa et al. 2016; Hasegawa and Arai 2017, 2018b, 2020a,b), information on wing length, migratory habit, clutch size, and incubation type was obtained from Turner and Rose (1994). We used the monograph (Turner and Rose 1994), because this publication provides detailed information on hirundines compared to general books on birds (although we referred to other publications as needed; see below). As before (e.g., Hasegawa and Arai 2018b), species were divided into migrant and nonmigrant species to account for the possible confounding factor, migration. We regarded species as migrant when they had breeding sites separate from their wintering sites (i.e., when they were summer visitors in a portion of their distribution range, which is shown in yellow in the color plates in Turner and Rose 1994). Whenever sex-specific values could be obtained, values for males were used, as before (e.g., Hasegawa and Arai 2018b). We also used values of nominate subspecies when subspecies differences were found.

We used the mean wing length as representative of species body size to account for the potential confounding effect of body size (Turner and Rose 1994; typos were corrected according to a personal communication from Angela Turner, i.e., the wing lengths of *H. angolensis* and *H. neoxena* are not 199 and 122 mm, but 119 and 112 mm, respectively). We used wing length as a proxy for body size, because wing length is tightly correlated with other measures of body size (e.g., body mass) and did not decrease the sample size (see detailed information in Hasegawa and Arai 2020a).

Data on clutch size were obtained from Hasegawa and Arai (2017) and were originally obtained from Turner and Rose (1994). Although clutch size and egg size are generally thought to be negatively correlated, a previous study found no detectable relationship between clutch size and egg size in hirundines, and thus we focused here on clutch size alone (Hasegawa and Arai 2017). When available, we also extracted information on the number of successful clutches (single vs. multiple) per season from Turner and Rose (1994). In this clade, hatching success is usually high, and thus brood sizes are similar to clutch sizes (note that fledging success can also be high; Turner and Rose 1994).

We also collected data on incubation type (female-only vs. biparental incubation) from the descriptions (e.g., “both sexes incubate,” “the female incubates alone”) in Turner and Rose (1994). When we were unable to obtain this information from Turner and Rose (1994), we searched for updates using del Hoyo et al. (2020). We adopted this dichotomous classification (Hasegawa and Arai 2020b), given the lack of information on fine-scale male incubation behaviors (e.g., the proportion of male incubation).

Finally, we extracted the adult-juvenile plumage discrepancy score, simply counting the notable discrepancies in plumage characteristics between adults and juveniles (i.e., the number of secondary sexual characteristics) in the notes for each plate of species in the text of Turner and Rose (1994). We focused on the notes of adults but not yearlings when the latter were available. For example, as a note on the barn swallow, Turner and Rose (1994) wrote “Juvenile: duller and browner, with rufous area paler; outer tail feathers are short,” and so we assigned a score of 3 for adult-juvenile plumage discrepancy for this species. This method may have underestimated the actual adult-juvenile plumage discrepancy, because these measures could not capture the degree of difference (e.g., how much duller, how short were the tails). However, this measure would still capture the number of different types of adult-juvenile plumage discrepancies in a predictable manner. Although in some bird species, immature birds have more colorful plumage than adults (e.g., Lyon et al. 1994), this is not the case for hirundines, since juveniles (i.e., post-fledging young) have duller, fainter, or more diffuse color patterns than adults (Turner and Rose 1994). An alternative method of comparing plumage between adults and juveniles using color plates (see Hasegawa and Arai 2018a for cuckoos) was not applicable in the current case, because plates showing juveniles are often absent in Turner and Rose (1994). Although the current method could not quantify ultraviolet reflection, ultraviolet reflection accompanies iridescent visible light reflection, at least in the barn swallow (e.g., Perrier et al. 2002), and thus the potential bias would be relatively small. We used male plumage characteristics, but qualitatively similar (and often the same) scores were obtained when we used female plumage characteristics, due to the small sexual dimorphism. Therefore, instead of studying male and female values separately, we recorded male-female plumage discrepancy, i.e., a measure of sexual dimorphism, which is easy to collect from the descriptions in the literature (because sex differences are described in the notes). Finally, absolute plumage ornamentation, measured as male plumage color score, was obtained from Dale et al. (2015). These scores, based on the nape, crown, forehead, throat, upper breast, and lower breast illustrated on the plate of the Handbook of the Birds of the World, reflect how ‘male-like’ a plumage is (note that other body regions were often not clearly shown and thus excluded; Dale et al. 2015). The data set is summarized in Table S1 (note that we list scientific names as reported in birdtree.org).

### Phylogenetic comparative analysis

We used a Bayesian phylogenetic mixed model with a normal error distribution to examine adult-juvenile plumage discrepancy. To account for potential confounding factors [body size, measured by log(wing length), and migratory habit], we included these variables along with clutch size in the model. To account for phylogenetic uncertainty (and phylogenetic inertia), we fit the models to each tree and used multimodel inference using 1000 alternative trees for swallows from birdtree.org (e.g., Garamszegi and Mundry 2014; Hasegawa and Arai 2017, 2018a,b). Similarly, to study the main and interaction terms of additional variables (i.e., clutch number and incubation type), we analyzed separate models using a subset of the data because of limited information (see tables for sample sizes). We derived mean coefficients, 95% credible intervals (CIs) based on the highest posterior density, and Markov chain Monte Carlo (MCMC)-based P-values, together with the phylogenetic signal (de Villemereuil and Nakagawa 2014). All analyses were conducted in R ver. 4.0.0 (R Core Team 2020) with the function “MCMCglmm” in the package “MCMCglmm” (ver. 2.29; Hadfield 2010) with the prior [G=list(G1=list(V=1, nu=0.02)), R=list(V=1, nu=0.02)]. We ran the analysis for 140000 iterations with a burn-in period of 60000 and a thinning interval of 80 for each tree.

## Results

### Abundance of young

There was no detectable linear relationship between the score of adult-juvenile plumage discrepancy and clutch size (P_MCMC_ = 0.67; data not shown). Likewise, there was no detectable linear relationship between the score of male-female plumage discrepancy and clutch size (P_MCMC_ = 0.57), and between male plumage color score and clutch size (P_MCMC_ = 0.90; data not shown). However, we found a quadratic relationship between clutch size and the score of adult-juvenile plumage discrepancy (Table 1): species with intermediate clutch sizes had the highest discrepancies in adult-juvenile plumage discrepancy while in species with small or large clutches, adults and juveniles resembled each other more closely (Fig. 2). No such quadratic pattern was found in male-female plumage discrepancy and male plumage color score (Table 1).

**Fig. 2.**
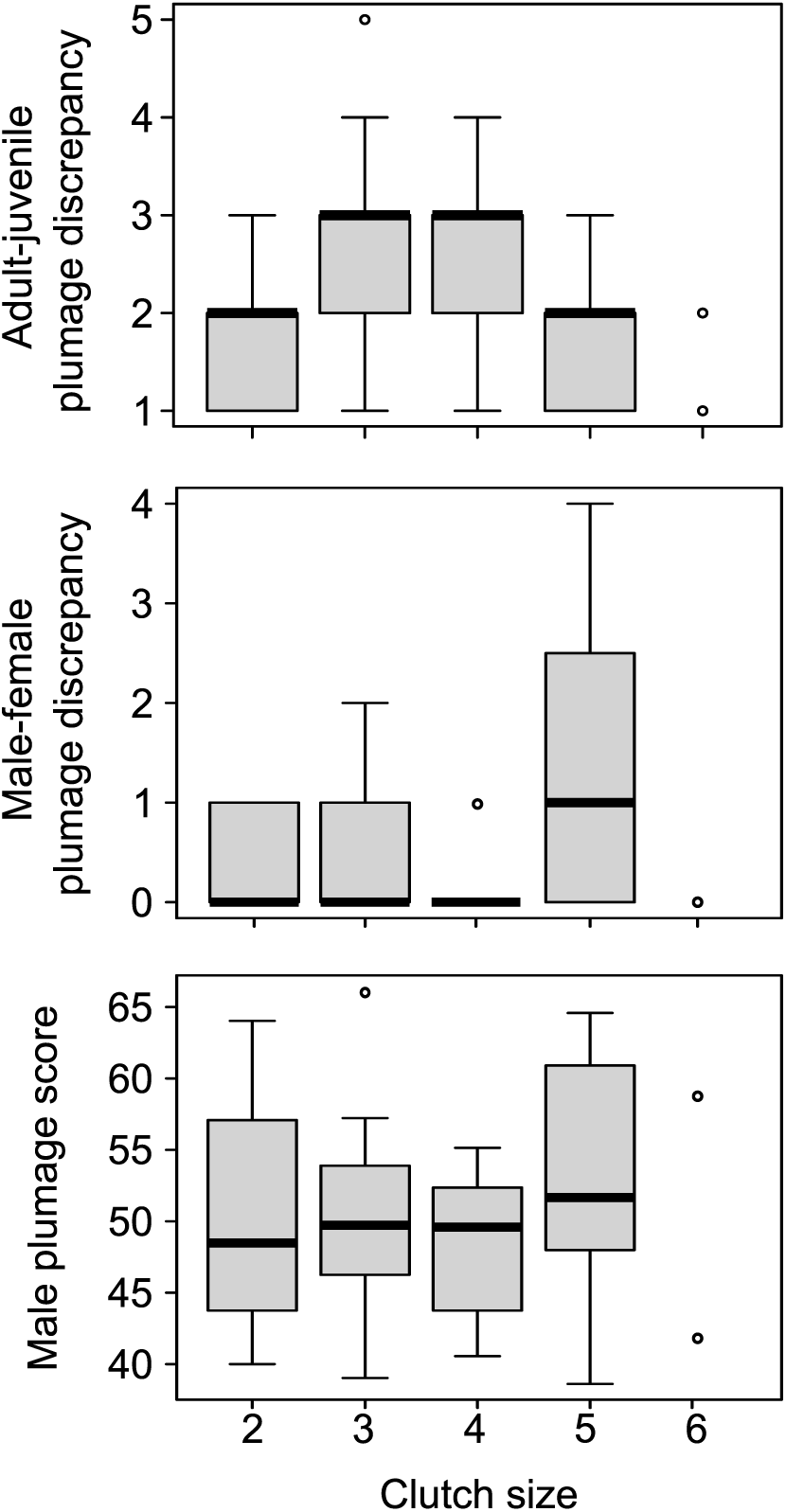
Boxplot of adult-juvenile plumage discrepancy (upper column), male-female plumage discrepancy (middle column), male plumage score (bottom column) in relation to mean clutch size in swallows and martins. The bars and boxes in each boxplot indicate medians and first and third quartiles, respectively. The whiskers range from the lowest to highest data points within 1.5 × the interquartile range of the lower and upper quartiles, respectively. Outliers are shown as circles (see Table 1 for detailed statistics). Number of species in each clutch size was 14, 26, 17, 8, 2 for clutch size 2, 3, 4, 5, 6, respectively. Note that response variable is natural number and thus each representative value of boxplot shape is also confined to natural number, limiting the shape of boxplot

**Table 1.** Multivariable Bayesian phylogenetic mixed model with a normal error distribution predicting (a) adult-juvenile plumage discrepancy, (b) male-female plumage discrepancy, (c) male plumage score, in relation to clutch size in swallows and martins (n = 67; Aves: Hirundinidae)

| Variable | Coefficient [95% CI] | P<sub>MCMC</sub> |
| --- | --- | --- |
| (a) adult-juvenile plumage discrepancy |  |  |
| log(wing length) | **0.30 [0.04, 0.55]** | **0.02** |
| Migration | -0.18 [-0.59, 0.24] | 0.40 |
| (Clutch size)<sup>2</sup> | **-0.22 [-0.38, -0.06]** | **<0.01** |
| Mean phylogenetic signal = 0.59 |  |  |
| (b) male-female plumage discrepancy after square-root transformation |  |  |
| log(wing length) | 0.07 [-0.09, 0.23] | 0.37 |
| Migration | 0.22 [-0.04, 0.48] | 0.09 |
| (Clutch size)<sup>2</sup> | -0.01 [-0.11, 0.08] | 0.83 |
| Mean phylogenetic signal = 0.58 |  |  |
| (c) male plumage score |  |  |
| log(wing length) | 0.46 [-1.36, 2.26] | 0.61 |
| Migration | 0.72 [-2.32, 3.75] | 0.64 |
| (Clutch size)<sup>2</sup> | 0.34 [-0.81, 1.47] | 0.55 |
| Mean phylogenetic signal = 0.69 |  |  |

Wing length was standardized to mean zero and unit variance after log-transformation.
Clutch size was centered to mean clutch size of 3.4.
Model-averaged coefficients and 95% credible intervals (CIs) are shown.
Significant results (i.e., P<sub>MCMC</sub> < 0.05) are indicated in bold.
The linear term of clutch size, if added to the model, was nonsignificant (a: P<sub>MCMC</sub> = 0.24; b: P<sub>MCMC</sub> = 0.60; c: P<sub>MCMC</sub> = 0.99) and thus removed from the models

When we included the number of successful clutches per season as an additional independent variable using a subset of the data (n = 39), we found that the score of adult-juvenile plumage discrepancy was significantly lower in species with multiple clutches than in species with a single clutch, although the sample size was roughly halved (Table 2; Fig. 3). The quadratic relationship between the score of adult-juvenile plumage discrepancy and clutch size remained significant in this model (Table 2), indicating that clutch size and number of clutches have a partially independent relationship with the score of adult-juvenile plumage discrepancy. When we examined male-female plumage discrepancy instead of adult-juvenile plumage discrepancy, the number of clutches was non-significant (Tables 2). This was also the case when we analyzed male plumage color score (Table 2).

**Fig. 3.**
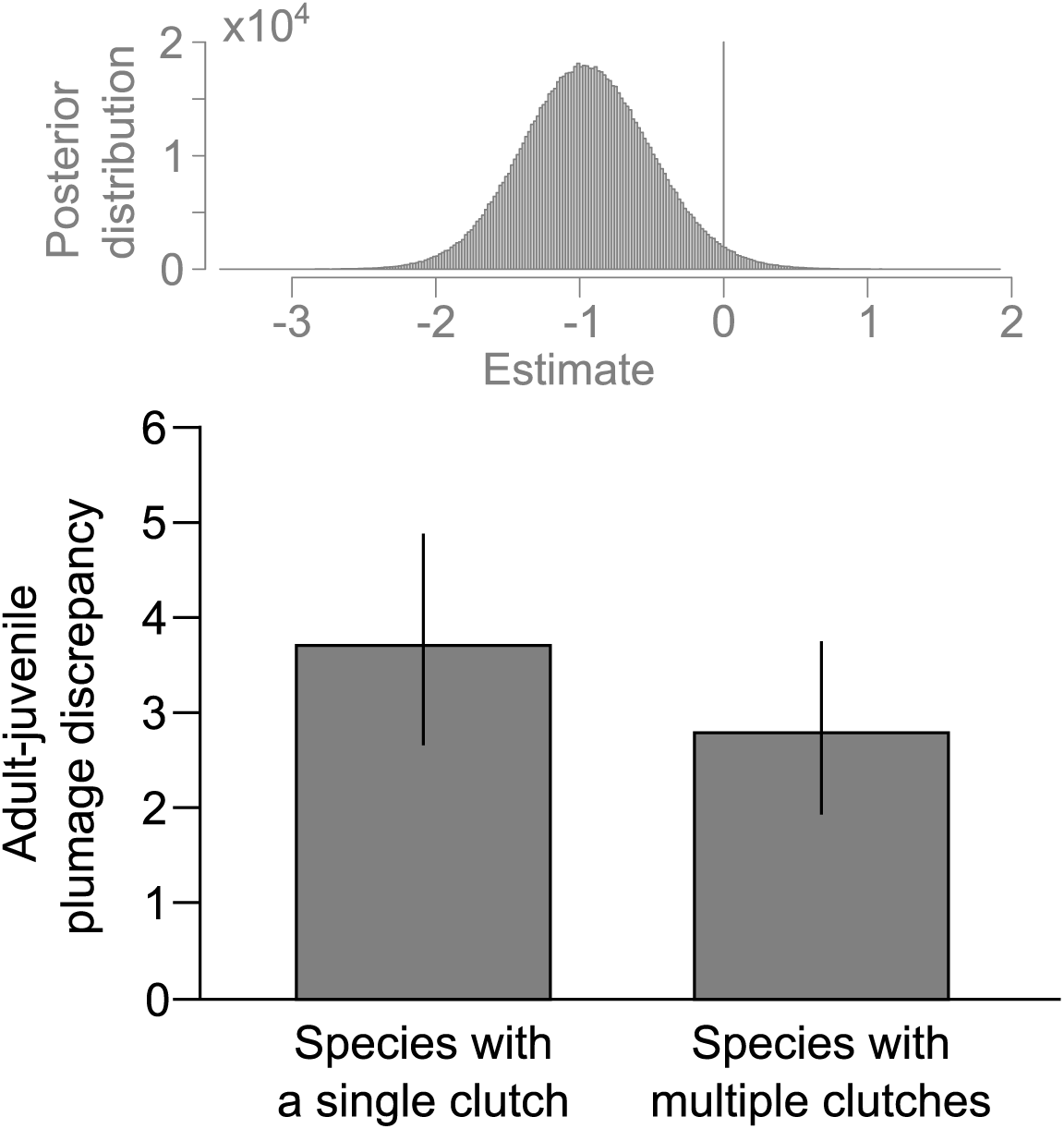
Predicted adult-juvenile plumage discrepancy in species with a single clutch and species with multiple clutches in swallows and martins. Means ± 95% CI are denoted. Above the boxplot, the posterior distribution of the estimated difference between the two groups sampled by MCMC chains across 1000 trees is shown. Detailed statistics are given in Table 3

**Table 2.**
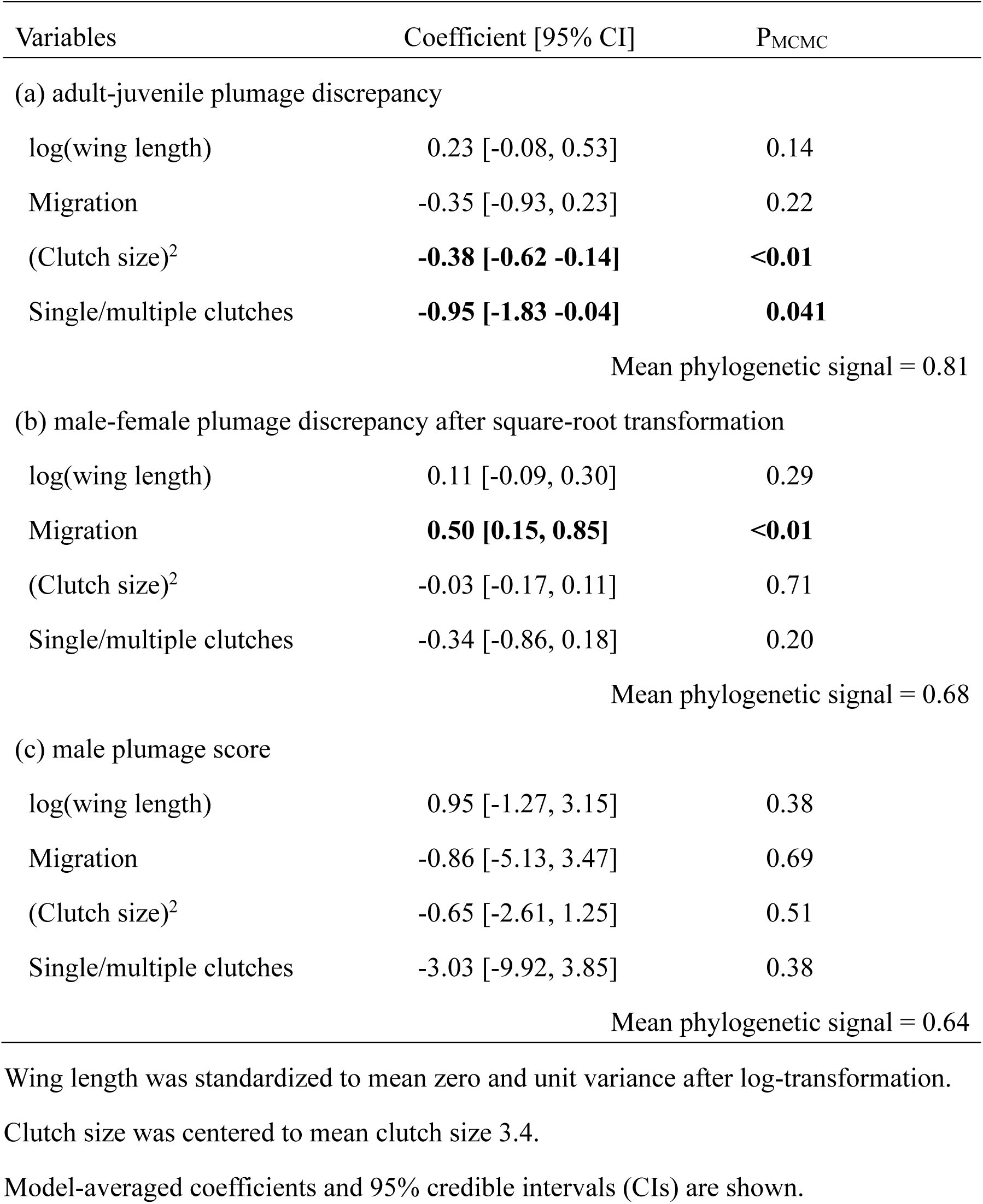
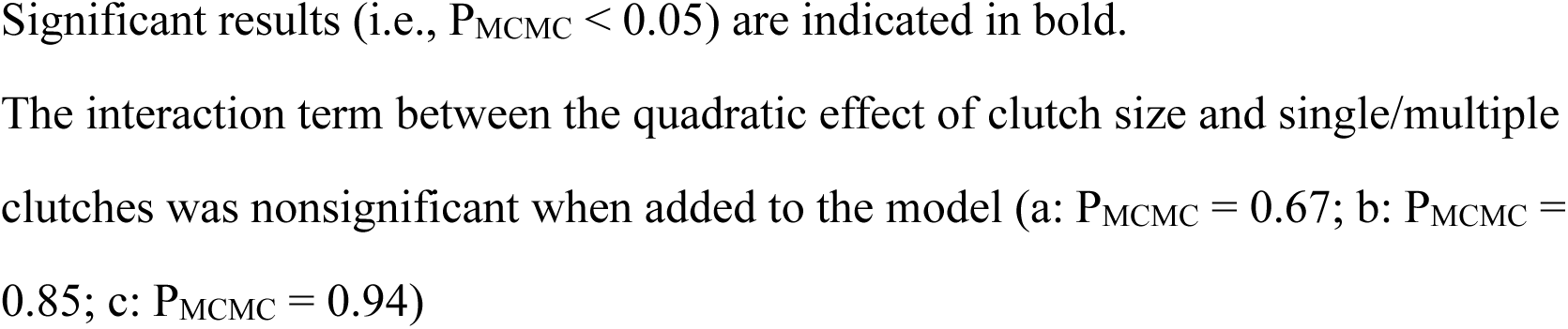
Multivariable Bayesian phylogenetic mixed model with a normal error distribution predicting (a) adult-juvenile plumage discrepancy, (b) male-female plumage discrepancy, (c) male plumage score, in relation to clutch size in swallows and martins (Aves: Hirundinidae) using species with information on number of clutches per year (n = 39)

| Variables | Coefficient [95% CI] | P<sub>MCMC</sub> |
| --- | --- | --- |
| (a) adult-juvenile plumage discrepancy |  |  |
| log(wing length) | 0.23 [-0.08, 0.53] | 0.14 |
| Migration | -0.35 [-0.93, 0.23] | 0.22 |
| (Clutch size)<sup>2</sup> | **-0.38 [-0.62 -0.14]** | **<0.01** |
| Single/multiple clutches | **-0.95 [-1.83 -0.04]** | **0.041** |
| Mean phylogenetic signal = 0.81 |  |  |
| (b) male-female plumage discrepancy after square-root transformation |  |  |
| log(wing length) | 0.11 [-0.09, 0.30] | 0.29 |
| Migration | **0.50 [0.15, 0.85]** | **<0.01** |
| (Clutch size)<sup>2</sup> | -0.03 [-0.17, 0.11] | 0.71 |
| Single/multiple clutches | -0.34 [-0.86, 0.18] | 0.20 |
| Mean phylogenetic signal = 0.68 |  |  |
| (c) male plumage score |  |  |
| log(wing length) | 0.95 [-1.27, 3.15] | 0.38 |
| Migration | -0.86 [-5.13, 3.47] | 0.69 |
| (Clutch size)<sup>2</sup> | -0.65 [-2.61, 1.25] | 0.51 |
| Single/multiple clutches | -3.03 [-9.92, 3.85] | 0.38 |
| Mean phylogenetic signal = 0.64 |  |  |

Wing length was standardized to mean zero and unit variance after log-transformation.
Clutch size was centered to mean clutch size 3.4.
Model-averaged coefficients and 95% credible intervals (CIs) are shown.
Significant results (i.e., $P_{\text{MCMC}} < 0.05$ ) are indicated in bold. The interaction term between the quadratic effect of clutch size and single/multiple clutches was nonsignificant when added to the model (a: $P_{\text{MCMC}} = 0.67$ ; b: $P_{\text{MCMC}} =$ $0.85$ ; c: $P_{\text{MCMC}} = 0.94$ )

**Table 3.**
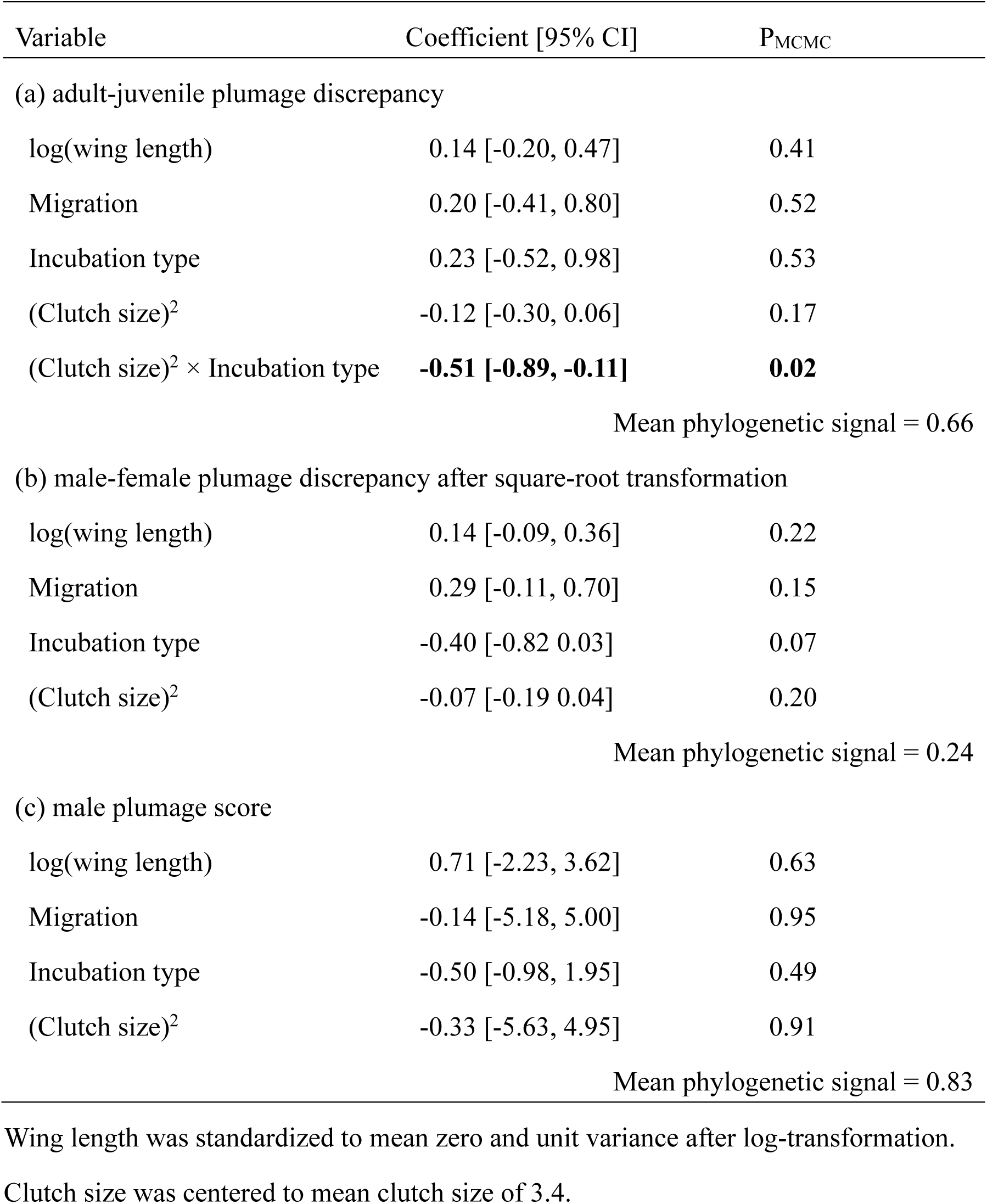
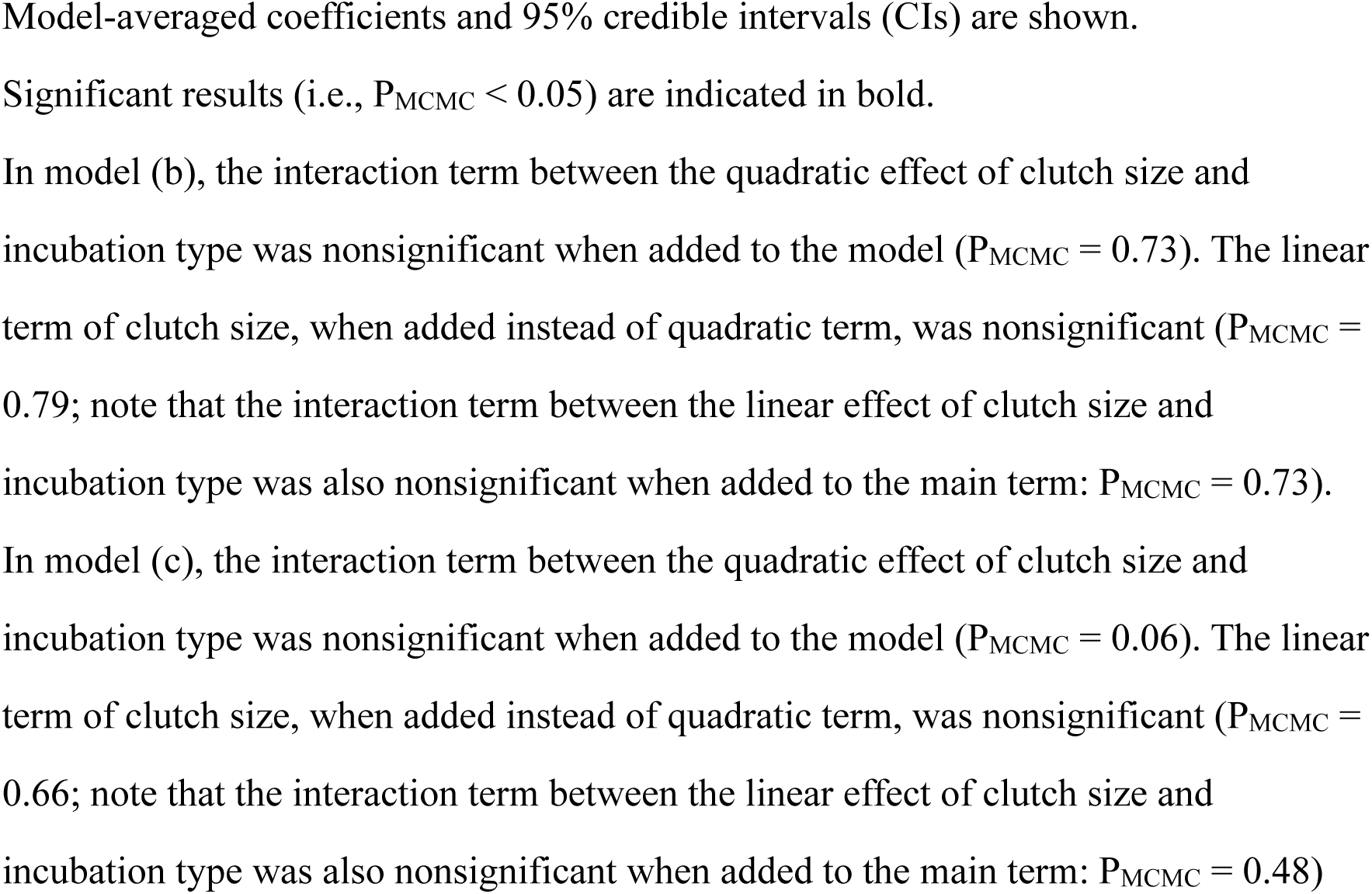
Multivariable Bayesian phylogenetic mixed model with a normal error distribution predicting (a) adult-juvenile plumage discrepancy, (b) male-female plumage discrepancy, (c) male plumage score, in relation to clutch size and incubation type (female-only vs. biparental incubation) in swallows and martins (n = 40; Aves: Hirundinidae)

| Variable | Coefficient [95% CI] | P<sub>MCMC</sub> |
| --- | --- | --- |
| (a) adult-juvenile plumage discrepancy |  |  |
| log(wing length) | 0.14 [-0.20, 0.47] | 0.41 |
| Migration | 0.20 [-0.41, 0.80] | 0.52 |
| Incubation type | 0.23 [-0.52, 0.98] | 0.53 |
| (Clutch size)<sup>2</sup> | -0.12 [-0.30, 0.06] | 0.17 |
| (Clutch size)<sup>2</sup> × Incubation type | **-0.51 [-0.89, -0.11]** | **0.02** |
| Mean phylogenetic signal = 0.66 |  |  |
| (b) male-female plumage discrepancy after square-root transformation |  |  |
| log(wing length) | 0.14 [-0.09, 0.36] | 0.22 |
| Migration | 0.29 [-0.11, 0.70] | 0.15 |
| Incubation type | -0.40 [-0.82, 0.03] | 0.07 |
| (Clutch size)<sup>2</sup> | -0.07 [-0.19, 0.04] | 0.20 |
| Mean phylogenetic signal = 0.24 |  |  |
| (c) male plumage score |  |  |
| log(wing length) | 0.71 [-2.23, 3.62] | 0.63 |
| Migration | -0.14 [-5.18, 5.00] | 0.95 |
| Incubation type | -0.50 [-0.98, 1.95] | 0.49 |
| (Clutch size)<sup>2</sup> | -0.33 [-5.63, 4.95] | 0.91 |
| Mean phylogenetic signal = 0.83 |  |  |

Wing length was standardized to mean zero and unit variance after log-transformation.
Clutch size was centered to mean clutch size of 3.4.
Model-averaged coefficients and 95% credible intervals (CIs) are shown. Significant results (i.e., $P_{MCMC} < 0.05$ ) are indicated in bold. In model (b), the interaction term between the quadratic effect of clutch size and incubation type was nonsignificant when added to the model ( $P_{MCMC} = 0.73$ ). The linear term of clutch size, when added instead of quadratic term, was nonsignificant ( $P_{MCMC} = 0.79$ ; note that the interaction term between the linear effect of clutch size and incubation type was also nonsignificant when added to the main term: $P_{MCMC} = 0.73$ ). In model (c), the interaction term between the quadratic effect of clutch size and incubation type was nonsignificant when added to the model ( $P_{MCMC} = 0.06$ ). The linear term of clutch size, when added instead of quadratic term, was nonsignificant ( $P_{MCMC} = 0.66$ ; note that the interaction term between the linear effect of clutch size and incubation type was also nonsignificant when added to the main term: $P_{MCMC} = 0.48$ ).

### Opportunities for extrapair paternity

When we included incubation type, as a measure of extrapair paternity, and its interaction term with clutch size, we found that the score of adult-juvenile plumage discrepancy was explained by the interaction between incubation type and the quadratic term of clutch size (Table 3). The score of adult-juvenile plumage discrepancy had a convex relationship with clutch size in species with biparental incubation (mean = - 0.63, 95% CI = -1.00, -0.25, P_MCMC_ < 0.01), whereas the quadratic term of clutch size on the score of adult-juvenile plumage discrepancy was weak and nonsignificant in species with female-only incubation (Table 3; Fig. 4). When we examined male-female plumage discrepancy instead of adult-juvenile plumage discrepancy, incubation type was marginal (Table 3). When we analyzed male plumage color score, none of the variables was significant (Table 3). Unfortunately, a meaningful analysis including clutch number and incubation type as additional independent variables could not be performed because only one species in this subset had a single clutch with biparental incubation (n = 32).

**Fig. 4.**
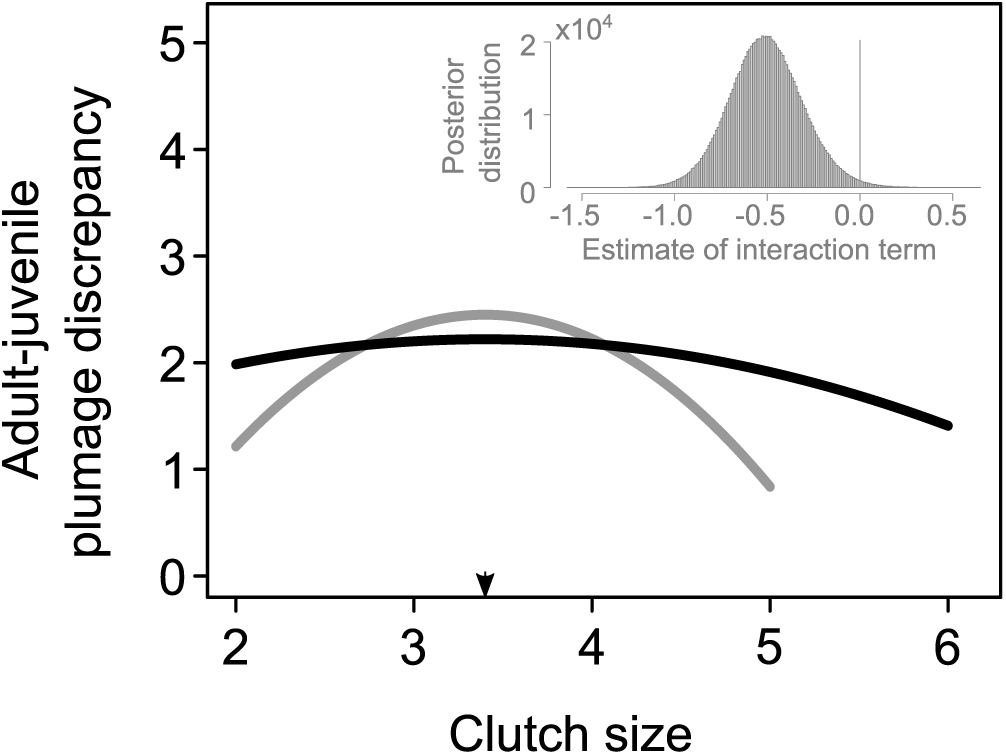
Estimated relationships between clutch size and adult-juvenile plumage discrepancy in species with female-only incubation (black line) and species with biparental incubation (gray line) after controlling for covariates in swallows and martins (note the clutch size ranges of 2–5 and 2–6 in species with female-only incubation and species with biparental incubation, respectively). The arrow indicates the mean clutch size (3.4). In the upper right corner, the posterior distribution of the estimated coefficient of interaction term sampled by MCMC chains across 1000 trees is shown (see Table 4 for detailed statistics)

## Discussion

The main finding of the current study is that adult-juvenile resemblance was enhanced in species with large clutches (i.e., species with many young) compared to species with mean clutch sizes (Table 1; Fig. 3), which is consistent with the juvenile-mimicry hypothesis. This finding is further reinforced by the enhanced adult-juvenile resemblance in multi-clutch species (Table 3), indicating that the presence of many young in general, rather than differential reproductive strategy (i.e., multiple small clutches vs. a single large clutch, here), explain the observed pattern. This aligns with the juvenile-mimicry hypothesis and supports the prediction that the mating period after juvenile production promotes juvenile-mimicry. Because male-female plumage discrepancy (i.e., a measure of costly plumage ornamentation) and male plumage color score (i.e. absolute male plumage appearance) showed no detectable relationship with the number of juveniles, sexual selection for secondary sexual characteristics alone or in combination with the cost of secondary sexual characteristics do not explain the observed pattern (see Introduction). For the same reason, intense selection for crypsis in species with many juveniles (e.g., to offset increased number of individuals and parental duty) is unlikely, because an absolute measure of ornamentation, i.e., male plumage score, did not vary systematically with the number of juveniles (though this possibility is inherently unlikely in this study system, i.e., hyper-aerial birds inhabit open space). In other words, male characteristics relative to those of young matters. The observed pattern is also inconsistent with the simple and widespread explanation of adult-juvenile resemblance, that is, adults in species with weak sexual selection retain juvenile-like plumage, though this “default” explanation (i.e., limited intensity of sexual selection) might account for high adult-juvenile resemblance in species with small clutches.

The differential relationships with clutch size between adult-juvenile plumage discrepancy and two other measures, namely male-female plumage discrepancy and male plumage color score, is not surprising, because the latter two measures were not significantly related to adult-juvenile plumage discrepancy (Univariable Bayesian phylogenetic mixed model with a normal error distribution: P_MCMC_ = 0.67 and 0.19; data not shown). This is further reinforced by the macroevolutionary pattern that male (and female) forked tails, a representative “costly” sexual trait known to reduce foraging ability due to aerodynamic cost, exhibit no detectable relationship with clutch size (Hasegawa and Arai 2017; also see Hasegawa 2024 for the cost of forked tails). Although macroevolutionary studies of sexual selection typically assume that all these measures reflect sexual selection for costly sexual traits, each measure might have evolved somewhat independently, serving distinct ecological functions (e.g., juvenile mimicry is specific to adult-juvenile resemblance; also see Tables 1–3 for male-female plumage discrepancy, but not others, had significant or marginal relationships with migratory habit and incubation type; note that sexual dimorphism reflects both inter-and intrasexual selection, meaning that some components can be independent of extrapair paternity).

The observed pattern, i.e., increased adult-juvenile similarity in large and multiple clutches, is consistent with the juvenile-mimicry hypothesis (see above); however, as a correlational study, the current study cannot completely exclude all possible explanations. For example, we did not distinguish the possibilities that juveniles evolved to resemble adults and that adults evolved to resemble juveniles (which cannot be separable from the current method; see Methods section), and thus juveniles might mimic adults in species with many young rather than adults mimic juveniles. This is, however, unlikely, because this possibility predicts the reverse pattern (i.e., a high number of juveniles prevents effective adult-mimicry and thus should weaken adult-juvenile resemblance). Or, juvenile-specific plumage characteristics (i.e., high adult-juvenile discrepancy) might be beneficial in species with an intermediate number of young. For example, adults might avoid agonistic interactions with typical juveniles when they have extrapair young (i.e., avoiding aggressions toward potential offspring). However, this is also unlikely as we found rather weaker effects of clutch size in species with high opportunities for extrapair mating (Table 3). For this reason, selection mediated through social interaction, rather than selection mediated through extrapair paternity, should be important to determine the adult-juvenile resemblance in relation to the number of young, as predicted by juvenile-mimicry hypothesis (see Introduction). Unfortunately, we could not exclude all other possibilities and confounding factors (e.g., sexual selection for secondary sexual characteristics combined with rearing cost of the traits might in theory make the same prediction for adult-juvenile discrepancy under certain situations, but overall pattern including male-female discrepancy and male plumage characteristics did not support them; see Introduction), which remain to be examined with experimental manipulation of adult-juvenile plumage discrepancies.

In summary, the current study supports the juvenile-mimicry hypothesis, explaining interspecific variation in adult-juvenile resemblance. Increased adult-juvenile resemblance in species with large clutches (and its interaction with incubation type) is difficult to explain by other hypotheses (see above). Although previous studies have dismissed the importance of juvenile-mimicry (e.g., see Rainey and Grether 2007 for a review of all studies of competitive mimicry, stating that juvenile mimicry “must be rare”), juvenile-mimicry might be a widespread phenomenon explaining adult-juvenile resemblance in species with parental care, particularly those with many young. Furthermore, selection for adult-juvenile resemblance counteracts sexual selection for (multiple) secondary sexual characteristics, potentially limiting ornament expression, or possibly, driving phenotypic diversification by reducing the number of secondary sexual characteristics (Figs. 3 & 4). Adult-juvenile resemblance would not always be a default state, as assumed in many studies, but would be an adaptive strategy in some situations, which should not be ignored in future studies. At the least, we should keep in mind that, although empirical studies have traditionally regarded relative plumage expression as a measure of sexual selection on male plumage characteristics (i.e., deviation from a default state; e.g., sexual plumage dimorphism, secondary sexual characteristics: see Introduction), the relative plumage expression (i.e., resemblance to other conspecific classes) can be the target of selection.

## Acknowledgments

We thank Dr Emi Arai, Dr Shumpei Kitamura and his lab members at Ishikawa Prefectural University for their kindest advices. We are grateful to Dr Angela Turner for her kindly support on the valuable information on swallows.

## Funding

MH was supported by the KAKENHI grant of the Japan Society for the Promotion of Science (JSPS, 19K06850).

## Author contribution

MH performed data analysis and wrote the manuscript.

## Data availability

The data sets supporting this article have been uploaded as part of electronic supplementary material, table S1.

## Declarations

### Conflict of interest

We have no competing interests

### Ethical approval

This comparative study does not include any treatments of animals, as all the information was gathered from literatures.

---

**Table S1.** Data set of the current study (n = 67)

| Latin name | Wing length (mm) | Migratory (1) or not (0) | Clutch size | Adult-juvenile plumage discrepancy* | Sexual plumage dimorphism† | Incubation period (days) | Nestling period (days) | No. clutches (1: single; 2: multiple) | Incubation type (1: female-only; 2: biparental) |
| --- | --- | --- | --- | --- | --- | --- | --- | --- | --- |
| *Alopochelidon fucata* | 100 | 1 | 4 | 1 | 0 | NA | NA | NA | NA |
| *Stelgidopteryx ruficollis* | 109 | 1 | 5 | 2 | 0 | 16 | 19 | 1 | 1 |
| *Stelgidopteryx serripennis* | 110 | 1 | 6 | 1 | 0 | 16 | 19 | 1 | 1 |
| *Tachycineta bicolor* | 119 | 1 | 5 | 3 | 1 | 14 | 19 | 1 | 1 |
| *Tachycineta albilinea* | 97 | 0 | 4 | 2 | 0 | 17 | 25 | 2 | NA |
| *Tachycineta albiventer* | 104 | 1 | 4 | 3 | 0 | NA | NA | NA | NA |
| *Tachycineta thalassina* | 122 | 1 | 5 | 2 | 1 | 14 | 23 | 1 | 1 |
| *Tachycineta leucorrhoa* | 115 | 1 | 6 | 2 | 0 | NA | NA | NA | 1 |
| *Tachycineta meyeri* | 110 | 1 | 5 | 1 | 0 | NA | NA | 2 | 2 |
| *Tachycineta cyaneoviridis* | 115 | 0 | 3 | 2 | 2 | NA | NA | NA | 1 |
| *Tachycineta euchrysea* | 106 | 0 | 3 | 3 | 2 | NA | NA | NA | NA |
| *Notiochelidon murina* | 111 | 0 | 2 | 3 | 0 | NA | NA | NA | NA |
| *Notiochelidon pileata* | 95 | 0 | 4 | 3 | 0 | NA | NA | NA | NA |
| *Pygochelidon cyanoleuca* | 94 | 1 | 4 | 3 | 0 | 15 | 26 | 2 | 2 |
| *Atticora fasciata* | 101 | 0 | 4 | 4 | 0 | NA | NA | NA | NA |
| *Atticora melanoleuca* | 93 | 0 | 3 | 3 | 0 | NA | NA | NA | NA |
| *Progne tapera* | 130 | 1 | 4 | 2 | 0 | 14 | 28 | NA | 1 |
| *Progne subis* | 145 | 1 | 5 | 2 | 4 | 16 | 28 | 1 | 1 |
| *Progne chalybea* | 131 | 1 | 5 | 2 | 3 | 17 | 26 | 1 | 1 |
| *Progne dominicensis* | 143 | 1 | 5 | 1 | 2 | NA | NA | 1 | NA |
| *Progne modesta* | 125 | 1 | 2 | 1 | 1 | NA | NA | NA | NA |
| *Riparia paludicola* | 104 | 0 | 3 | 2 | 0 | 12 | 20 | NA | 2 |
| *Riparia riparia* | 107 | 1 | 5 | 1 | 0 | 14 | 22 | 2 | 2 |
| *Riparia cincta* | 130 | 1 | 3 | 2 | 0 | NA | NA | NA | NA |
| *Psalidoprocne fuliginosa* | 104 | 0 | 2 | 1 | 0 | NA | NA | NA | NA |
| *Psalidoprocne albiceps* | 102 | 1 | 3 | 2 | 1 | NA | NA | NA | 2 |
| *Psalidoprocne*<br>*pristoptera* | 106 | 1 | 2 | 2 | 0 | 14 | 25 | NA | 1 |
| *Psalidoprocne*<br>*obscura* | 96 | 0 | 2 | 2 | 1 | NA | NA | NA | NA |
| *Psalidoprocne*<br>*nitens* | 93 | 0 | 2 | 1 | 0 | NA | NA | NA | NA |
| *Cheramoeca*<br>*leucosterna* | 102 | 0 | 5 | 3 | 0 | NA | NA | 1 | NA |
| *Pseudhirundo*<br>*griseopyga* | 97 | 0 | 3 | 3 | 0 | NA | NA | NA | NA |
| *Phedina*<br>*borbonica* | 116 | 0 | 2 | 1 | 0 | NA | NA | NA | 1 |
| *Phedina*<br>*brazzae* | 100 | 0 | 3 | 2 | 0 | NA | NA | NA | NA |
| *Hirundo*<br>*rupestris* | 130 | 1 | 3 | 1 | 0 | 15 | 25 | 2 | 1 |
| *Hirundo*<br>*fuligula* | 129 | 0 | 2 | 1 | 0 | 17 | 26 | 2 | 2 |
| *Hirundo*<br>*concolor* | 106 | 0 | 3 | 2 | 0 | NA | NA | 2 | 2 |
| *Hirundo*<br>*rustica* | 124 | 1 | 4 | 3 | 1 | 15 | 20 | 2 | 1 |
| *Hirundo*<br>*lucida* | 111 | 0 | 3 | 2 | 0 | NA | NA | NA | NA |
| *Hirundo*<br>*angolensis* | 119 | 0 | 3 | 3 | 0 | NA | NA | 2 | NA |
| *Hirundo*<br>*tahitica* | 105 | 0 | 2 | 3 | 0 | 16 | 20 | 2 | 1 |
| *Hirundo*<br>*neoxena* | 112 | 0 | 4 | 3 | 0 | 16 | 20 | 2 | 1 |
| *Hirundo albigularis* | 128 | 1 | 3 | 3 | 1 | 15 | 20 | 2 | NA |
| *Hirundo aethiopica* | 106 | 0 | 4 | 3 | 0 | 14 | 23 | 2 | 1 |
| *Hirundo smithii* | 110 | 1 | 3 | 2 | 1 | 14 | 19 | 2 | 1 |
| *Hirundo nigrata* | 106 | 0 | 3 | 2 | 0 | 15 | 17 | 2 | 1 |
| *Hirundo leucosoma* | 99 | 0 | 4 | 2 | 0 | NA | NA | NA | NA |
| *Hirundo dimidiata* | 102 | 1 | 2 | 2 | 1 | 16 | 21 | 2 | 1 |
| *Hirundo atrocaerulea* | 113 | 1 | 3 | 3 | 1 | 15 | 22 | 2 | 1 |
| *Hirundo nigrorufa* | 112 | 1 | 3 | 3 | 1 | NA | NA | NA | NA |
| *Hirundo cucullata* | 125 | 1 | 3 | 4 | 1 | 18 | 26 | 2 | 1 |
| *Hirundo abyssinica* | 106 | 1 | 3 | 4 | 1 | 15 | 18 | 2 | 1 |
| *Hirundo semirufa* | 132 | 1 | 3 | 4 | 0 | NA | NA | 2 | 1 |
| *Hirundo senegalensis* | 144 | 0 | 3 | 5 | 0 | NA | NA | 2 | NA |
| *Hirundo daurica* | 124 | 1 | 3 | 4 | 0 | 14 | 24 | 2 | 2 |
| *Hirundo striolata* | 124 | 0 | 4 | 4 | 0 | NA | NA | NA | NA |
| *Hirundo preussi* | 95 | 0 | 2 | 3 | 0 | NA | NA | NA | NA |
| *Hirundo rufigula* | 97 | 1 | 2 | 2 | 0 | NA | NA | NA | 2 |
| *Hirundo spilodera* | 111 | 1 | 2 | 2 | 0 | 14 | 24 | 2 | 2 |
| *Hirundo fuliginosa* | 88 | 0 | 2 | 1 | 1 | NA | NA | NA | NA |
| *Petrochelidon pyrrhonota* | 109 | 1 | 3 | 4 | 0 | 13 | 24 | 1 | 2 |
| *Petrochelidon fulva* | 102 | 1 | 4 | 2 | 0 | 15 | 24 | 2 | 2 |
| *Hirundo fluviicola* | 91 | 1 | 3 | 3 | 0 | NA | NA | 2 | 2 |
| *Hirundo ariel* | 91 | 1 | 4 | 2 | 0 | NA | NA | 2 | 2 |
| *Hirundo nigricans* | 107 | 1 | 4 | 3 | 0 | NA | NA | 2 | NA |
| *Delichon urbicum* | 110 | 1 | 4 | 2 | 0 | 15 | 27 | 2 | 2 |
| *Delichon dasypus* | 108 | 1 | 3 | 2 | 0 | NA | NA | 2 | 2 |
| *Delichon nipalense* | 95 | 0 | 3 | 2 | 0 | NA | NA | 2 | 2 |

NA indicates Not available.
\*, †: See text for detailed explanation

## Notes

### Competing Interest Statement

The authors have declared no competing interest.

